# Whole genome sequencing and variant discovery in 344 global grasspea (*Lathyrus sativus* L.) lines

**DOI:** 10.64898/2026.06.05.730453

**Authors:** Miriam Schreiber, Jasmine Staples, Peter M.F. Emmrich, Anne Edwards, Cathie Martin, Micha Bayer, Sebastian Raubach, Benjamin Kilian, Paul D. Shaw

## Abstract

The rapid expansion of genomic data resources for major crops is opening new options for crop improvement, while resources for most underutilised crops lag behind, risking a widening gap in crop improvement. One of these underutilised crops is grasspea (*Lathyrus sativus*), an ancient crop with modern cultivation centred on South Asia and Ethiopia. We conducted whole genome shotgun sequencing on a global collection of 344 grasspea lines, producing over 152 billion reads. Following variant discovery and filtering we created a single nucleotide polymorphism (SNP) marker set of over 1.5 million SNPs. This is a resource of major significance for this crop which can help unlock its breeding potential through marker development and the identification of genes controlling agronomically important traits.

## Background & Summary

Grasspea (*Lathyrus sativus L.*) has been cultivated for at least 8000 years, spreading across northern Africa, Asia and Europe^1^. Nowadays, cultivation of grasspea is centred on South Asia and the highlands of Ethiopia, but also European countries such as Portugal. Grasspea is diploid and has seven chromosomes, with a large genome size of approx. 6.3 Gbp. A high-quality genome assembly of the genotype LS007 has recently been published^2^. Grasspea is predominantly self-pollinated, but can outcross^3^, resulting in variable levels of heterozygosity in the population. Cultivar breeding is restricted by lack of knowledge about the genetic control of relevant traits, such as production of the neurotoxin β-L-oxalyl-diaminopropionic acid (β-ODAP) or disease resistance. While there have been recent studies of disease resistance against powdery mildew^4^, fusarium^5^ and rust^6^, the shortage of genetic resources restricts the identification of causal genes and marker development for breeding. Existing studies have relied on simple sequence repeats (SSRs)^7^ or a small subset of single nucleotide polymorphisms (SNPs)^5^ for genome wide association studies (GWAS). While genetic resources have been increasing in recent years, grasspea is still very much under-resourced in comparison to major crops. A search of the European Nucleotide Archive for by taxonomy ID, revealed 151,022 sequencing datasets for wheat, 139,849 for rice and 8352 for pea in comparison to 261 sequencing datasets for grasspea (accessed: 05/06/2024). Here, we are expanding the knowledge base for grasspea by providing 344 whole genome shotgun sequencing datasets which have been mapped to the reference genome and used for variant calling to develop a robust SNP dataset. This resource will enable downstream analyses like GWAS and marker development to support breeding.

## Methods

### Grasspea plant material and tissue sampling

Grasspea seeds were obtained from the ICARDA (International Center for Agricultural Research in the Dry Areas) Genebank in Terbol, Lebanon together with accession passport data, including original collection sites and dates and alternative names (http://grs.icarda.org/; Supplemental Table 1). This collection includes lines and data previously shared by USDA National Plant Germplasm System, the Leibniz Institute of Plant Genetics and Crop Plant Research (IPK), the Bangladesh Agricultural Research Institute and the John Innes Centre (JIC). All lines were regenerated at Terbol, Lebanon. Plants were grown in a glasshouse in John Innes Multipurpose and Grit: (90% peat, 10% grit, 4 kg/m3 Dolomitic Limestone, 0.75 kg/m3 PG mix, 1.5kg/m3 osmocote bloom). Seeds were started in 24 cell trays then transplanted to F11 pots (one plant/pot) and grown in a glasshouse at a day temperature of 20°C and a night temperature of 16°C, and a photoperiod of 16hrs. Plants were watered from below on ebb and flood automatic benches, once per day for newly sown and very young plants, increasing to twice per day.

### Flower colour scoring

Plants for flower colour scoring were grown in an unlit, unheated polytunnel at the JIC with planting in March 2023. Seeds were directly planted into John Innes Multipurpose and Grit in 5L pots (5 plants of the same lines/pot). Scoring for flower colour took place between the 14^th^ of June and 3^rd^ July, due to differences in flowering time. Photographs of at least one representative flower from each plant were taken, and scored as blue-violet, white, mostly white flowers with blue colouration only on the wings and the centre of the standard petal, or pink. Any unusual flower colours were noted, and for any lines with variation in flower colour between plants, this was noted, captured in the image and the line excluded from further analysis.

### Sequencing

Genomic DNA was extracted from 40 mg fresh 3-week-old leaf tissue using Octopure sbeadex− magnetic bead technology (JIC Genotyping platform). Samples were diluted to 20 ng/ul and 200 ul were supplied to Novogene for library preparation. Whole genome shotgun sequencing (WGS) was conducted by Novogene using the Illumina NovaSeq 6000 sequencing platform, generating 150bp paired end reads.

### Bioinformatics

### Mapping and variant calling

Whole genome shotgun Illumina sequencing reads were aligned to the grasspea genome (GCA_963859935.3)^2^ using bwa-mem (v0.7.17)^8^ with default parameters. Samtools (v1.6)^9,10^was used to mark duplicates in the alignment files, followed by filtering with sambamba (v1.0, with parameters --paired-pairs --mapping_quality >= 60)^11^ to remove multi-mapped reads and reads with low confidence in the mapping location. For variant calling only reads with a mapping quality of 60 were kept, avoiding unreliably mapped reads. Variant calling was conducted with the Genome Analysis Toolkit (GATK) v4.2.6^12^ by using the GATK HaplotypeCaller module on each BAM file individually to produce a genomic variant calling file (gVCF). The 344 gVCF files from all samples were combined using GATK CombineGVCF on a per chromosome basis. The read mapping, filtering of the BAM files and variant calling steps were combined into a Snakemake workflow^13,14^ which can be found on GitHub (https://github.com/SchreiberM/BOLD_Grasspea_SNPSet). Genotyping of samples was carried out with GATK GenotypeGVCF in 20Mbp genomic intervals^15,16^ on the combined gVCF file. This resulted in 285,816,635 raw variants, with 265,767,427 of these SNPs the remainder being small insertions and deletions.

### Filtering

For better compatibility, we aimed to extract a set of 5,651 previously published SNP markers^5^ from the raw variant dataset for inclusion in our final set of markers (see below). The previous published SNP markers were searched against the genome using BLAST^17^ with at least 90% identity and only unique hits considered. The SNP position was extracted and a total of 2,606 were present in the raw variant dataset. These were filtered to keep only bi-allelic SNP markers (vcftools --recode-INFO-all --min-alleles 2 --max-alleles 2--remove-indels --recode; vcftools v0.1.16^18^).

The remaining variants from the raw VCF file were first filtered for a mean maximum depth of 25 reads, a mean minimum depth of 2 reads (bcftools view --include ‘INFO/DP>670 && INFO/DP<8375’; bcftools v1.9)^10^ and at least one homozygous genotype (bcftools view --include ‘COUNT(GT=‘\”hom’\”)>1’) to avoid variants which are completely heterozygous across all genotypes. Further filtering was conducted with vcftools to keep only bi-allelic SNPs with a minor allele frequency of 0.1, a minimum variant quality of 100 and no missing data (vcftools --recode-INFO-all --maf 0.01 --min-alleles 2 --max-alleles 2 --max-missing 1 --remove-indels --minQ 100 --recode).

This resulted in a filtered dataset of 10,448,963 SNPs. A marker set of over 10 million SNPs is challenging for most downstream analyses and computing resources, and thus this dataset was further reduced in size by pruning SNPs using linkage disequilibrium (LD) and by prioritising SNPs in genes over those in intergenic spaces as it has been shown that intergenic SNPs are less likely to be associated with complex traits^19^.

Therefore the 10 million SNPs were first split into a gene space and an intergenic dataset (vcftools --exclude-bed GeneSpace.bed; vcftools --bed GeneSpace.bed). The GeneSpace.bed file was generated from the gff file corresponding to the published genome (GCA_963859935.3)^2^ by extracting the coordinates for the type ‘\”gene’\”. The gene space VCF file contained 1,190,089 SNPs while the intergenic space file contained 9,258,874 SNPs. Plink (v.1.90b6.21)^20^ was used to filter for LD, the gene file with --allow-extra-chr --indep-pairwise 10000 kb 1 0.9 and the intergenic file with --allow-extra-chr --indep-pairwise 10000 kb 1 0.7, reducing the dataset to 723,744 SNPs in the gene space and 1,613,235 in the intergenic regions. Finally, all three datasets -- the previously published SNPs, the gene space SNPs and the intergenic SNPs -- were combined into the final SNP marker set.

### Variant annotation

We used SnpEff^21^ to add genomic variant annotations to the final SNP marker set by creating a SnpEff database from the published genome (GCA_963859935.3).

### Quality control

Multiple statistics were generated to report on the quality of the final SNP marker set. A multi-dimensional scaling plot was created using plink --cluster --allow-extra-chr --mds-plot 3. Nucleotide diversity was calculated over 1 Mbp intervals using vcftools --window-pi 1000000 --window-pi-step 1000000. The number of heterozygous SNPs at each position was extracted from the output of vcftools --hardy.

### Genome-wide association study

For the genome-wide association study (GWAS) the program GAPIT (version 3)^22^ was run in R (v4.3)^23^ by providing the phenotype results (flower colour in Supplemental Table 3) and the final SNP marker set with default parameters for the following four models: MLM^24^, MLMM^25^, BLINK^26^ and FarmCPU^27^.

### Data Records

The raw data files and the marker dataset are available on request.

### Technical Validation

Whole genome shotgun sequencing of 344 grasspea lines resulted in over 152 billion raw reads corresponding to over 22 terabasepairs (Supplemental Table 1). Read mapping showed that on average 89.22% of the reads were properly mapped and paired, and average genome coverage of the raw unfiltered mapping results was 13.1x. As grasspea has highly repetitive regions in its genome^2^, a high proportion of reads (between 25.42% to 32.54%) showed a low mapping quality of MQ0 (Supplemental Table 1). Variant calling was done using GATK4, calling variants across all samples simultaneously. Over 285 million variants were identified and of those over 265 million were SNPs. Variant sets of this size are unsuitable for most downstream analyses, and thus further filtering was required for the dataset to be useable for applications such as GWAS or marker development for breeding. Therefore, we conducted stringent quality filtering and filtering by LD. A subset of previously published SNPs^5^ were carried over and combined with our filtered SNP set. In the last step the dataset was filtered for a minor allele frequency of >0.05, an established cut-off for most GWAS analyses. This resulted in the final marker dataset of 1,049,647 SNPs.

#### Multi-dimensional scaling plots

As a first quality check we generated a multi-dimensional scaling plot to establish whether the SNP markers were a good representation of the population and clustered the lines based on collection sites of the accessions from which they were derived as has been previously shown^5^. The first three dimensions were calculated for the marker dataset. As shown in Figure 1, the first dimension separates Asian and European lines. The second dimension splits off the East African lines, most of which originate from Ethiopia. The third dimension separates the Asian lines into two different clusters: Central Asian, characterised by lines from Afghanistan and Tajikistan, and South Asian, characterised by lines from Nepal, Bangladesh and Pakistan. This is comparable to the principal component analysis conducted on the much smaller SNP dataset from Sampaio et al^5^ which showed a similar picture, with the first principal component differentiating between European and Asian lines and the second component separating the Ethiopian lines.

**Figure 1.**
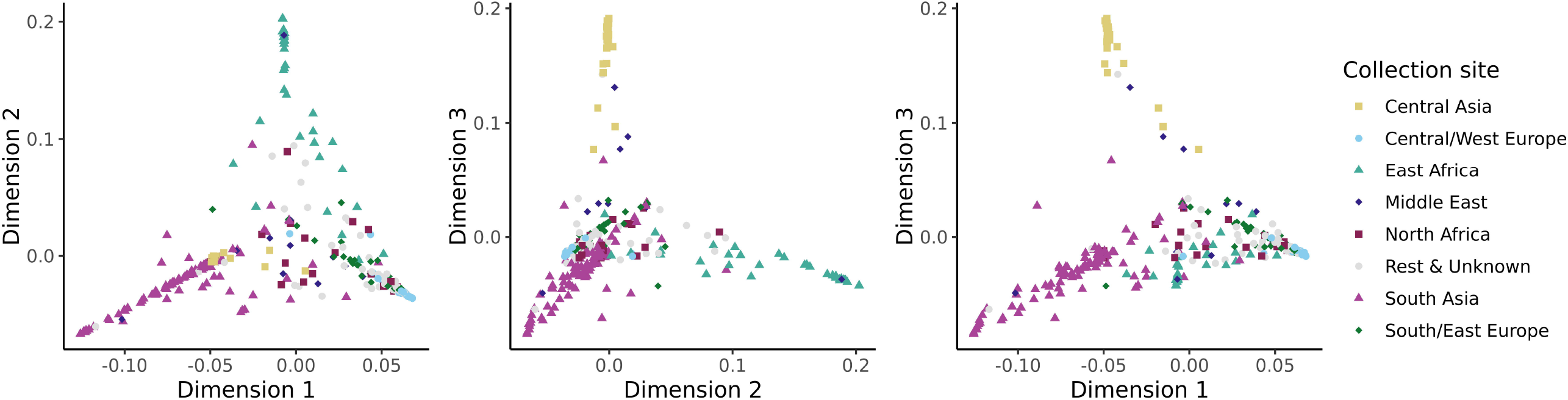
Multidimensional scaling plot of the 344 grasspea lines based on the SNP marker dataset. Colours are based on the collection site of each original accession (Supplemental Table 1).

#### Variant annotation using SnpEff

The 1,049,647 SNPs were annotated with 1,510,736 entries in the SnpEff output. The discrepancy in numbers is due to some SNPs being associated with multiple annotations, for example due to simultaneously being upstream of one gene and downstream of another gene, or in genes overlapping on opposite strands. Most of the annotations (53.36%) were in intergenic regions, but 46.64% were found within or close to genes (1kbp upstream and downstream), with 8.48% located within exons (Table 1).

**Table 1.** SnpEff results of the SNP marker set.

| Type (alphabetical order) | Count | Percentage |
| --- | --- | --- |
| DOWNSTREAM | 237,187 | 15.7% |
| EXON | 128,040 | 8.48% |
| INTERGENIC | 806,114 | 53.36% |
| INTRON | 138,710 | 9.18% |
| SPLICE_SITE_ACCEPTOR | 32 | 0.00% |
| SPLICE_SITE_DONOR | 44 | 0.00% |
| SPLICE_SITE_REGION | 6,863 | 0.45% |
| UPSTREAM | 193,746 | 12.83% |

The results from the nucleotide diversity and distribution of heterozygous SNPs were compared to the chromosomal annotation from the previous genome publication^2^. The gene density (Figure 2a) along the seven chromosomes of grasspea was reflected in the SNP distribution (Figure 2b) with a broadly comparable pattern. Higher SNP and gene density was evident towards the telomeric ends of the chromosomes, with pericentromeric regions showing an inverse trend. Pericentromeric regions, which have been shown to be rich in repetitive elements and low in gene content^2^, were less diverse while telomeric regions with higher gene content also had higher nucleotide diversity (Figure 2 c). A number of exceptions to this were observed in the centromeric regions of chromosomes Lschr1, Lschr2 and Lschr6, which overlapped with clusters of SNPs with higher heterozygosity (Figure 2 d). This finding may be related to the highly diverse metapolycentric centromeres found in grasspea^28^. As grasspea is capable of outcrossing, heterozygous SNPs are to be expected, and these showed an even distribution across chromosomes, except for the regions listed above. Most of the SNPs (80.7%, 846,860) had less than 10% heterozygosity. Most lines had a heterozygosity to homozygosity ratio of <1, meaning they were homozygous at most SNP locations, but 17.2% (59/344) of the lines had a >1 ratio (Supplemental Table 2).

**Figure 2.**
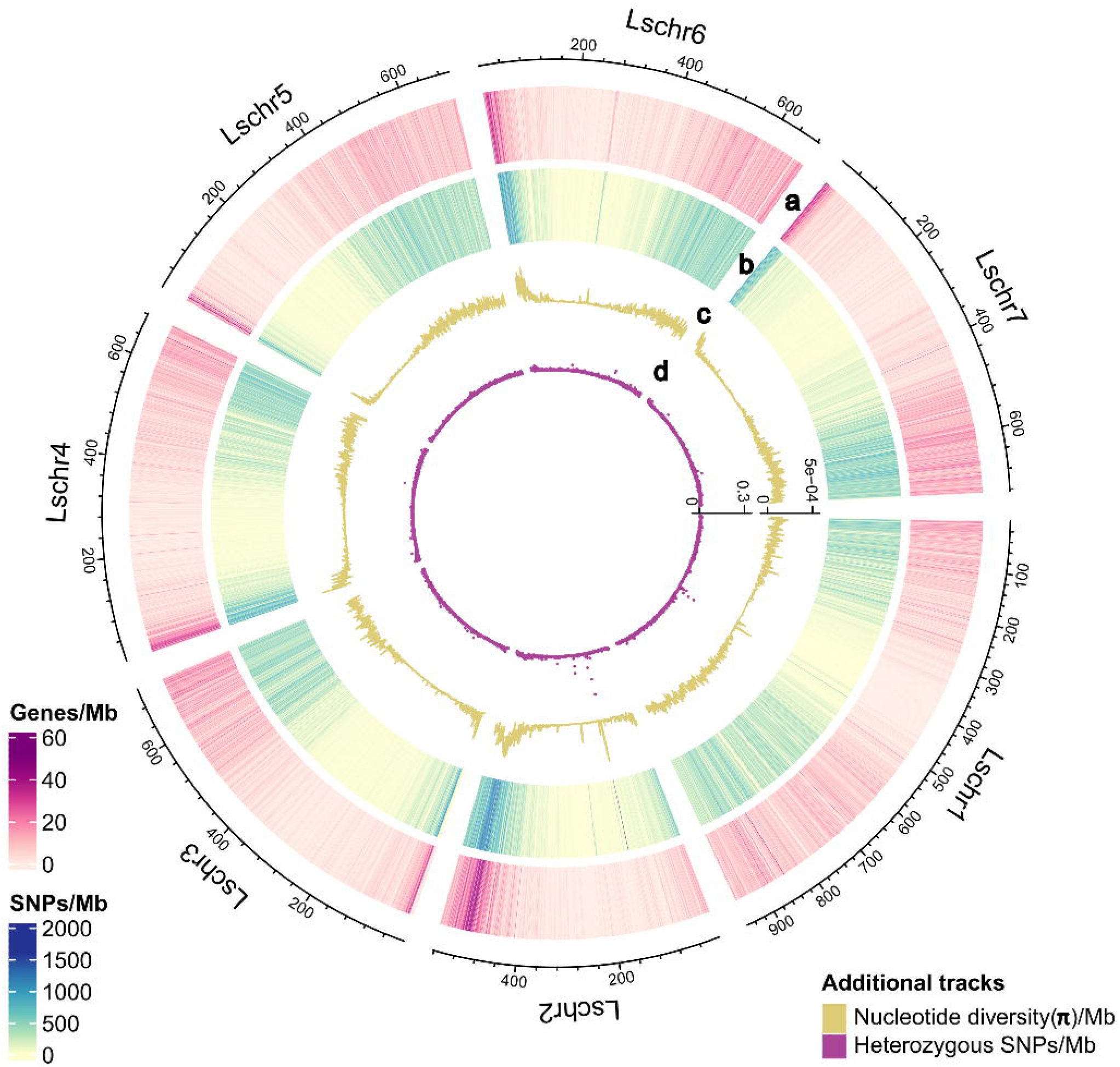
Overview statistics of the SNP marker set. (a) Gene density in number of genes per Mb, (b) SNP density in number of SNPs per Mb, (c) Nucleotide diversity (rr) per Mb, (d) number of heterozygous SNPs per Mb.

### Usage Notes

#### This section is optional

One downstream use case for the marker dataset is as input into genome wide association studies (GWAS). As an illustration of the potential of our marker set for GWAS, we conducted an analysis of flower pigmentation, which was first studied in peas by Gregor Mendel and has since been shown to be controlled by a bHLH transcription factor at the dominant A locus^29^. Grasspea lines were grown in a polytunnel, and the colour of the mature flowers was recorded. Flower colour was classified as blue-violet, white, or a distinct phenotype of mostly white flowers with blue colouration only on the wings and the centre of the standard petal. Plants with flowers that could not be clearly assigned to one of the three categories mentioned above, including a small subset of pink-flowered lines, were recorded as NA for the purposes of this analysis (Supplemental Table 3). GWAS revealed a clear peak on Lschr6 for flower colour (Figure 3) for all four models used with the GAPIT software. These markers colocalise with the grasspea homologue of the bHLH gene (Lsat_6G0002460400; gene ID LATHSAT_LOCUS24531; protein ID CAK8571908) at the pea A locus, with the highest significant SNP (OZ012740.1_492299957) located in an exon. Further downstream analysis could be done to identify and verify if the underlying gene corresponds to the A gene. This example highlights the potential of our marker dataset for use in identifying genes underlying agronomically relevant traits.

**Figure 3.**
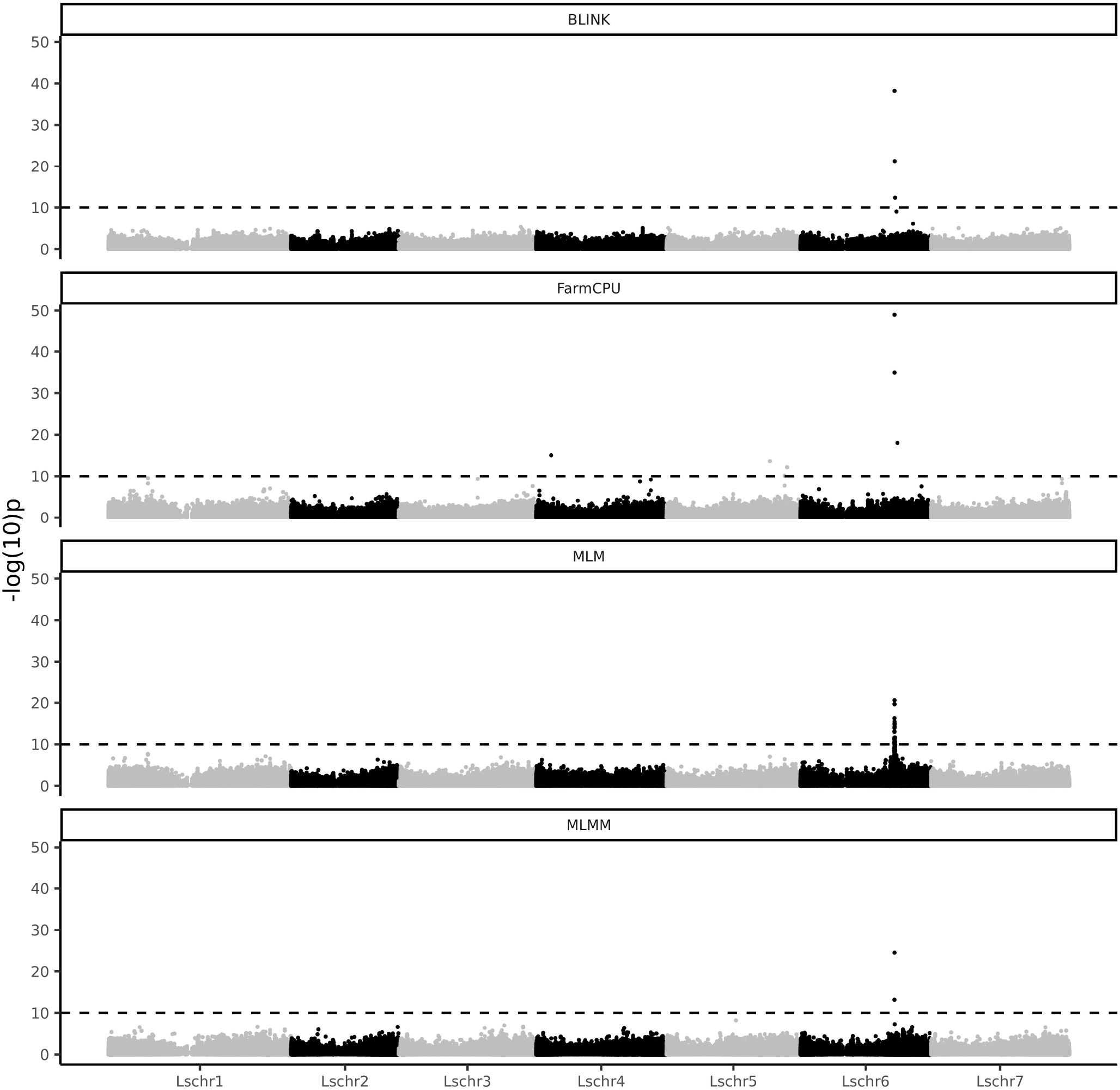
GWAS analysis for flower colour in the grasspea collection using the final SNP marker set and four different GAPIT models.

## Code Availability

All code is either available through the GitHub repository: https://github.com/SchreiberM/BOLD_Grasspea_SNPSet or the parameters are described in detail in the methods section.

## Acknowledgements

The authors acknowledge Research Computing at the James Hutton Institute for providing computational resources and technical support for the ‘\”UK’s Crop Diversity Bioinformatics HPC’\” (BBSRC grants BB/S019669/1 and BB/X019683/1), use of which has contributed to the results reported within this paper.

This work is part of the Crop Trust BOLD project, which is generously funded by the Government of Norway. BOLD (Biodiversity for Opportunities, Livelihoods and Development) is a 10-year project to strengthen food and nutrition security worldwide by supporting the conservation and use of crop diversity (grant number: QZA-20/0154). For more information visit https://bold.croptrust.org. This work was also supported by the project ‘Safeguarding Crop Diversity for Food Security: Pre-breeding complemented with Innovative Finance’, supported by the Templeton World Charity Foundation, Inc. (Grant ID TWCF0400).

## Author contributions

M.S. did the bioinformatics analysis; M.S., P.E., M.B. wrote the manuscript; J.S. and A.E. did the phenotyping; P.E., C.M., B.K., P.S. contributed to conceptual guidance and project development.

